# Oxytocin facilitates empathic- and self-embarrassment ratings by attenuating amygdala and anterior insula responses

**DOI:** 10.1101/343616

**Authors:** YaYuan Geng, Weihua Zhao, Feng Zhou, Xiaole Ma, Shuxia Yao, Benjamin Becker, Keith M. Kendrick

## Abstract

The hypothalamic neuropeptide oxytocin has been reported to enhance emotional empathy in association with reduced amygdala activation, although to date studies have not investigated empathy for individuals expressing self-conscious, moral emotions which engage mentalizing as well as emotion processing networks. In the current randomized, double-blind placebo controlled functional MRI experiment on 70 male and female subjects we have therefore investigated the effects of intranasal oxytocin (40 IU) on behavioral and neural responses to embarrassment experienced by others or by self. Results showed that oxytocin significantly increased ratings of both empathic and self-embarrassment and concomitantly decreased skin conductance response and activation in the right amygdala and insula but not in the medial prefrontal cortex. The amygdala effects of oxytocin were associated with the magnitude of the skin conductance response and trait anxiety scores. Overall our results demonstrate that oxytocin increases ratings of self- and other embarrassment and that this is associated with reduced physiological arousal and activity in neural circuitry involved in emotional arousal. The neural effects of oxytocin are also stronger in individuals with high trait anxiety suggesting that it may particularly reduce their anxiety in embarrassing situations.

## Introduction

Our ability to empathize with others is a core feature influencing our social behavior through allowing us to understand both what others are thinking and feeling and thereby promoting our social interactions with them. As such, impaired ability to empathize with others is often a core feature of disorders where social communication and interactions are dysfunctional, such as autism spectrum disorder (Lombardo et al., 2007), depression (Tully et al., 2016) and psychopathy (Ali et al., 2009).

Even though empathy has been extensively investigated, its sub-components and associated neural mechanisms are still not fully established (Moya-Albiol et al., 2010). The most prevalent view considers empathy as a multidimensional construct including both cognitive (identifying emotions expressed by another person) and emotional components (being aroused by or feeling the same emotion expressed by another person (Shamay-Tsoory, 2011). Meta-analytic data has suggested that a network including left orbital frontal cortex, left anterior mid-cingulate cortex, left anterior insula and left dorsal medial thalamus is involved in cognitive-evaluative empathy, whereas another network including the dorsal anterior cingulate cortex, bilateral anterior insula, right dorsal medial thalamus and midbrain is involved in affective-perceptual empathy (Fan et al., 2011). Studies that directly compared the two empathy components additionally proposed that the inferior frontal gyrus is essential for emotional empathy, whereas the superior and middle frontal gyrus and the orbital gyrus are specifically engaged in cognitive empathy (Preston et al., 2007). However, the derived empathy network can differ dependent upon the paradigm used (Moya-Albiol et al., 2010) thus making it important to establish a core network that is maintained across different paradigms.

The hypothalamic neuropeptide oxytocin (OXT) has been implicated in a number of crucial aspects of social cognition and emotional bonds (Striepens et al., 2011; Kendrick et al., 2017) and importantly studies have reported that it can enhance components of empathy (Hubble et al., 2017; Hurlemann et al., 2010; Geng et al. preprint). The most consistent findings have been in studies using paradigms that distinguish cognitive from emotional empathy components. For example, in male Caucasian subjects OXT was found to enhance both direct and indirect aspects of emotional empathy, but not cognitive empathy *per se*, in the Multifaceted Empathy Task (MET) (Hurlemann et al., 2012). Urbach-Wiethe disorder patients with selective and complete bilateral amygdala-lesions also exhibited deficits in emotional but not cognitive empathy in the MET suggesting that the emotional empathy enhancing effects of OXT might be mediated by the amygdala (Hurlemann et al., 2012). We recently confirmed these behavioral findings in Chinese male, as well as female subjects using a Chinese version of the MET and demonstrated that OXT effects on emotional empathy were associated with decreased amygdala activity and increased physiological arousal (Geng et al., preprint). Finally, another study using dynamic, empathy-inducing video clips in which protagonists expressed sadness, happiness, pain or fear demonstrated that OXT exerted no effects on cognitive empathy but selectively enhanced emotional empathy for fear (Hubble et al., 2017). Thus, OXT may particularly enhance emotional empathy, although in the context of the strong modulatory influence of personal and social contextual factors on the specific effects of OXT (Kendrick et al., 2017; Olff et al., 2013) it is important to extend these observations using across other paradigms and contexts where empathic responses are evoked.

Some initial support for OXT also influencing aspects of cognitive empathy have been reported using the reading the mind in the eyes test (RMET) where participants identify from visual cues restricted to the eye regions which of four different complex emotions is being experienced by subjects. Thus, in one study OXT was reported to increase accuracy, particularly for more difficult items (Domes et al., 2007). However, subsequent studies found that OXT either only increased RMET accuracy for difficult items in individuals with low empathy scores (Feeser et al., 2015) or generally only in individuals with lower socially proficiency associated with higher levels of maternal love withdrawal (Riem et al., 2014). Furthermore, a final comprehensive study failed to observe any effects of OXT on RMET performance even taking into account item difficulty, gender and valence and subject traits (Radke et al., 2015). Thus, evidence for OXT enhancing cognitive empathy using the RMET paradigm remains equivocal.

In the current double-blind, placebo-controlled study we therefore investigated whether OXT would enhance empathy in social situations involving more complex moral, “self-conscious”, emotions such as embarrassment. Embarrassment is primarily a self-reflection and self-evaluative process which serves as a way to help humans adapt their behavior to social norms by punishing non-compliance to such norms with a negative emotional state (Melchers et al., 2015). The neural pathways involved in embarrassment include both those controlling mentalizing and emotional arousal (Krach et al., 2016). Oxytocin has previously been shown to modulate activity in both of these networks in the context of processing self-referential information (mPFC – (Zhao et al., 2016; Zhao et al., 2015) and emotional stimuli (amygdala and anterior insula – Scheele et al., 2014; Kirsch et al., 2005; Gao et al., 2016, Yao et al., 2017).

We can experience empathic-embarrassment (EE) towards others we see in embarrassing situations irrespective of whether they are familiar or not (Miller, 1987) and so in the current paradigm we investigated OXT effects on behavioral and neural responses to pictures depicting others in embarrassing situations. Additionally, we also asked subjects to imagine their own feelings if they experienced a similar situation themselves (i.e. self-embarrassment – SE). We reasoned that the latter self-related context would involve a stronger mentalizing component, although both contexts should have strong arousal components. We hypothesized that OXT would increase both EE and SE ratings and that in both cases this would be associated with reduced amygdala and insula activation. In view of the greater self-orientation and mentalizing component in the SE condition we additionally hypothesized that OXT would increase mPFC activation in this condition. We have previously shown that some behavioral and neural effects of OXT on empathy are associated with increased physiological arousal and modulated by autistic traits (Geng et al., preprint). The neural effects of embarrassment are also modulated by anxiety (Mueller-Prinzler et al., 2015) and autism (Krach et al., 2016). Thus, in the current study we also investigated the effects of OXT on physiological arousal as assessed by electrodermal activity and associations of its effects with autism and anxiety traits.

## Methods and Materials

### Participants

A total of n = 70 participants were enrolled in the present randomized double-blind, placebo-controlled between-subject experiment. Participants were randomly assigned to receive OXT (40 International Units, IU) or placebo (PLC) intranasal treatment resulting in 35 participants (female n=17) in the OXT treatment group and 35 (female n=15) in the PLC treatment group. Both groups were of comparable age (mean ± STD, OXT: 22.03 ± 2.15; PLC: 21.86 ± 1.97; p = 0.73, t(70) = -0.35), education (OXT: 16.09 ± 1.65; PLC: 15.89 ± 1.64; p = 0.61, t(70) = -0.51), and gender distribution (*χ*^2^ (1) = 0.23, p = 0.63). All participants reported having no past / current physical, psychiatric or neurological disorders or regular / current use of medication or tobacco. Subjects were required to refrain from nicotine, alcohol or caffeine intake for at least 12 hours before the experiment. None of the female participants were taking oral contraceptives or were in their menstrual period. The distribution of females estimated to be in their follicular or luteal phases did not differ significantly between the groups (*χ*^2^ = 0.06, *p* = 1.00). Written informed consent was obtained from each participant before the experiment. The study was approved by the ethical committee of the University of Electronic Science and Technology of China, and all procedures were in accordance with the latest revision of the declaration of Helsinki.

### Experimental paradigm

40 pictures depicting male or female protagonists (Chinese, 20 male, 20 female protagonists) in everyday embarrassing situations were evaluated by an independent sample (n = 29, 14 females) prior to the experiment to balance mean ratings for empathic embarrassment (EE – how embarrassed do you feel for the person in the picture?) and self-embarrassment (SE – if you were in the same situation, how embarrassed would you feel?) in response to the pictures and the gender of the protagonist (EE: females, 6.54 ± 1.14, males, 6.65 ± 0.89; SE: females, 7.91 ± 0.55, males, 7.58 ± 0.67). No significant differences were found with regard to both EE and SE ratings between male and female participants (EE, p = 0. 77, t = -0.30; SE, p = 0. 17, t = 1.41). For the fMRI experiment stimuli were presented in a mixed block/event related design during four subsequent runs. Each run containing one block of EE and one block of SE trials, with ten trials of stimuli presented during each block. Each block started with a 3 seconds(s) cue presentation indicating whether the subject was required to rate EE or SE. Within each block stimuli were presented for 3 seconds, followed by a jittered low-level baseline during which a fixation cross was presented for 4s (3-5s). After each stimulus, subjects were given 5s to rate EE or SE using a 1-9 Likert rating scale (1 = not at all, 9 = very strong) followed by another 5s (4-6s) jittered inter-trial interval. An ABBA block-design was used to counterbalance the order of conditions (see Figure 1 for paradigm details).

**Figure 1.**
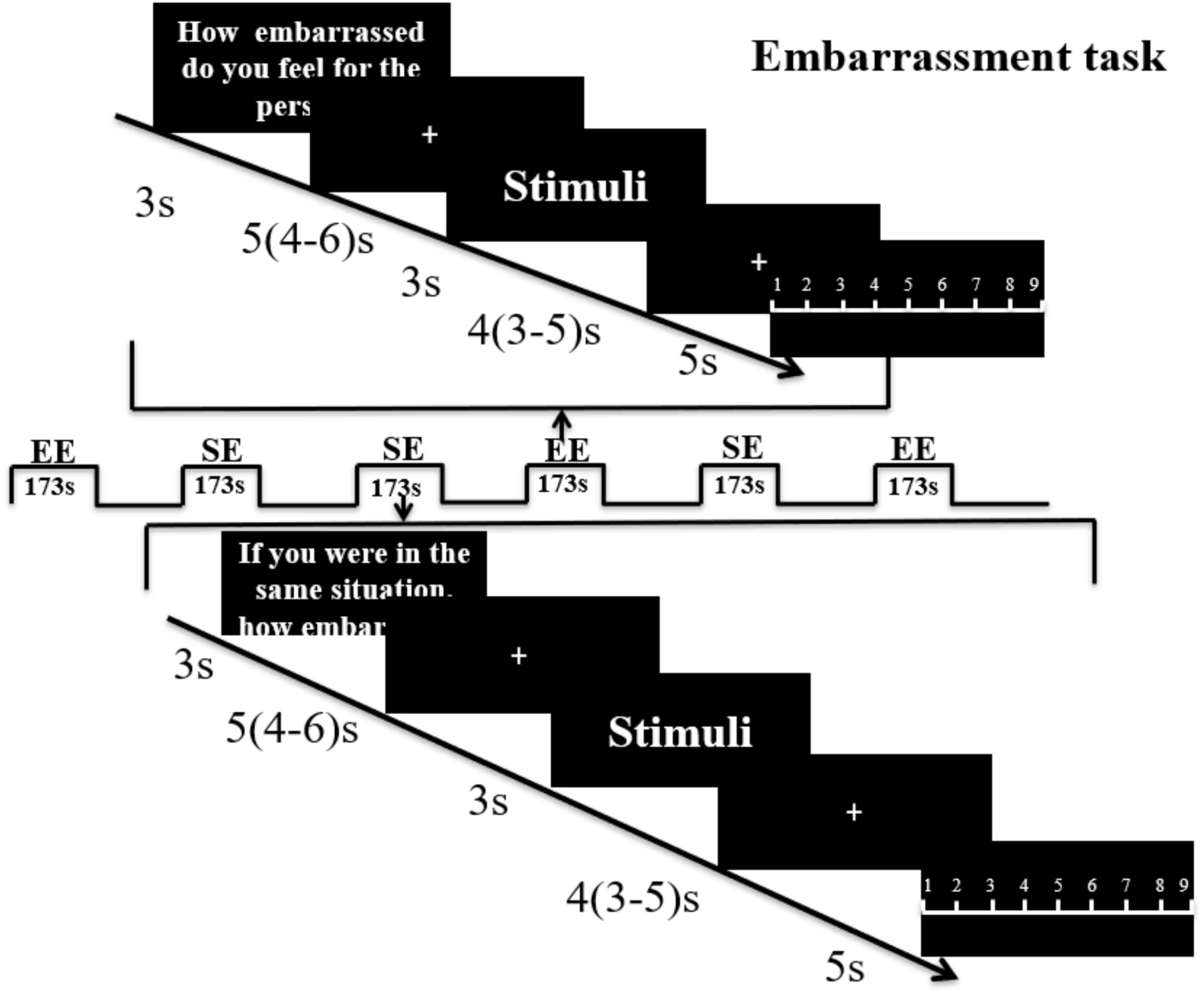
Paradigm for the embarrassment task. Subjects were first shown an instruction for 3s to indicate whether it was an empathic embarrassment (EE) or self-embarrassment (SE) trial followed by a jittered fixation cross (5s mean duration with 4-6 s range. The picture stimulus was then presented for 3s, followed by another fixation jittered mean duration of 4s (range 3 to 5s). A rating slide (1-9) was then shown for 5s. Each run included one EE block and one SE block of 10 stimuli and the run order was counterbalanced using an ABBA design.

### Procedure

To control for potential confounders, all participants completed a test battery of Chinese versions of mood and trait questionnaires before intranasal administration: Beck’s Depression Inventory (BDI) (Beck, 1961), Emotional Intelligence Scale (Wleis-C) (Wong et al., 2002), Empathy Quotient (EQ) (Baron-Cohen et al., 2004), Liebowitz Social Anxiety Scale (LSAS) (Liebowitz, 1987) and Positive and Negative Affect Scale (PANAS) (Watson et al., 1988). Based on previous research reporting that individual variations in autism and anxiety influence (1) embarrassment-associated neural activity (Miller et al., 1987; Mueller-Pinzler et al., 2015), as well as (2) effects of OXT on embarrassment-related functional domains (e.g. emotional arousal, empathy and self-appraisal) and neural activity in embarrassment-related regions, including the insula (Geng et al., preprint; Alvares et al., 2012; Scheele et al., 2014; Schumacher et al., 2018; Bartz et al., 2010), levels of anxiety and autism were assessed in the present sample. To this end, participants additionally completed the State Trait Anxiety Inventory (STAI) (Spielberger et al., 1970) and Autism Spectrum Quotient (ASQ) (Baron-Cohen et al., 2001). Next, subjects self-administered intranasal spray (either 40IU of OXT or PLC lacking the neuropeptide, both supplied by the Sichuan Meike Pharmaceutical Co., Ltd, Sichuan, China). The PLC spray had identical packing and ingredients as the OXT spray (sodium chloride and glycerine, minus the peptide). In accordance with previous recommendations for the intranasal administration of OXT in humans (Kendrick et al., 2017; Gustaella et al., 2013), the experimental paradigm started 45 minutes after treatment. In post-experiment interviews participants were unable to guess better than chance which treatment they had received (34 subjects guessed correctly; *χ*^2^ = 0.22, *p =* 0.64), confirming successful blinding.

During the experiment electrodermal activity was also measured to assess skin conductance responses (SCR) to the stimuli as an index of autonomic sympathetic activity (Stern, 2001) using the same approach as previously described (Geng et al., preprint). For the SCR data an event-related analysis approach was employed focusing on SCR responses associated with the presentation of SE and EE stimuli (procedures for preprocessing and event-related analysis of the SCR data were identical to our previous study, details provided in (Geng et al. preprint).

### fMRI acquisition

MRI data was acquired using a GE (General Electric Medical System, Milwaukee, WI, USA) 3T Discovery 750 MRI system with a standard head coil. fMRI time series were acquired using a T2*-weighted echo planar imaging pulse sequence (repetition time, 2000ms; echo time, 30ms; slices, 39; thickness, 3.4 mm; gap, 0.6 mm; field of view, 240 × 240 mm^2^; resolution, 64 × 64; flip angle, 90°). Additionally, a high resolution T1-weighted structural image was acquired using a 3D spoiled gradient recalled (SPGR) sequence (repetition time, 6 ms; echo time, 2ms; flip angle 9°; field of view, 256 × 256 mm^2^; acquisition matrix, 256 × 256; thickness, 1 mm without gap) to exclude subjects with apparent brain pathologies and to improve normalization of the fMRI data.

### fMRI data processing

fMRI data were analyzed using SPM12 (Wellcome Trust Center of Neuroimaging, University College London, London, United Kingdom). The first five volumes were discarded from further analyses and images were realigned to the first image to correct for head motion using a six-parameter rigid body algorithm and unwarping. Tissue segmentation, bias-correction and skull-stripping were performed for the high-resolution structural images. Functional images were further corrected for slice-acquisition time differences, co-registered to the anatomical scan and subsequently spatially normalized to the standard Montreal Neurological Institute (MNI) template. The normalized functional volumes were written out at a 3mm × 3mm × 3mm voxel size and were finally smoothed with an 8mm full-width-at-half-maximum (FWHM) isotropic Gaussian kernel.

On the first level, separate event-related regressors for the EE and SE conditions were included as main regressors of interest. Additionally, separate regressors for the rating phases, the cue phase and the six head motion parameters were included. The regressors were convolved with the standard hemodynamic response function (HRF). To evaluate the sex-dependent effects of OXT on embarrassment an ANOVA including treatment (OXT, PLC), sex (male, female) as between-subject factors and embarrassment type (EE, SE) as a within-subject factor was conducted on the second level.

Based on previous studies indicating that regions involved in arousal (amygdala, anterior insula, AI) and mentalizing (mPFC) neurally underpin embarrassment processing a region-of-interest (ROI) analysis specifically focused on these regions. The amygdala was structurally defined based on masks from the Automated Anatomical Labeling (AAL) atlas (Tzourio-Mayzoyer et al., 2002). Given the size and functional heterogeneity of these regions, the ROIs for the mPFC and anterior insula (specifically dorsal AI, dAI) were defined using 6mm spheres centered at peak coordinates of these regions reported in a previous study examining the neural basis of embarrassment (Mueller-Pinzler et al., 2015). For the five a priori defined ROIs the first eigenvariate was extracted using Marsbar (Brett et al., 2002). The individual activity estimates were subsequently subjected to mixed ANOVAs with the between-subject factors treatment (OXT, PLC) and sex (male, female) and the within-subject factor embarrassment type (EE and SE) in SPSS (Statistical Package for the Social Sciences, Version 22). Multiple comparisons for the ROI analysis were controlled for by applying False Discovery Rate (FDR) correction. To explore effects in regions beyond the predefined ROIs an exploratory whole brain analysis was conducted in SPM using cluster-level Family-Wise Error (FWE) correction for multiple comparisons. Significance level for the corrected p-values was p < .05.

## Results

No significant trait and mood differences were observed between participants in the OXT and PLC groups (see Supplementary Table S1).

### Behavioral and SCR results

An ANOVA analysis with treatment and sex as between-subject factors, embarrassment context as a within-subject factor and embarrassment rating as dependent variable, revealed a significant treatment main effect (PLC: 6.45 ± 0.11, OXT: 6.77 ± 0.11, F (1,66) = 4.30, p = 0.04, η^2^_p_ = 0.06) with OXT enhancing embarrassment ratings in both contexts (Figure 2). No significant interaction effects involving treatment were found (Sex * Treatment, F (1,66) = 2.20, p = 0. 14, η^2^_p_ = 0.03; Sex * Treatment * embarrassment context, F (1,66) = 0.09, p = 0.76, η^2^_p_ = 0.001).

**Figure 2.**
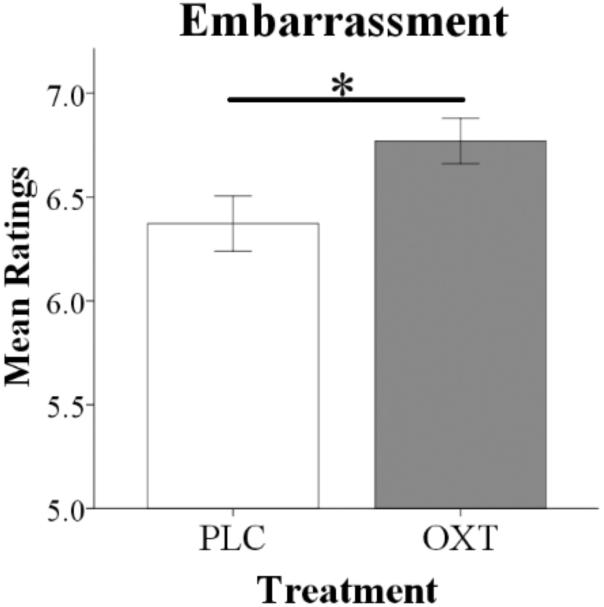
Oxytocin significantly increased overall embarrassment ratings in behavioral tests. Histograms show mean ratings for EE and SE trials combined (* p<0.05)

As a result of scanner-induced noise SCR data from n = 12 subjects had to be excluded leading to a final sample size of n = 58 subjects for the SCR analysis (n = 33, OXT; n = 25, PLC). An ANOVA analysis including the same factors as for the behavioral analysis and SCR response as dependent variable revealed a significant main effect of treatment (F (1,54) = 7.35, p = 0.009, η^2^_p_ = 0.12) with OXT decreasing the SCR magnitude in both EE and SE conditions (see Figure 3).

**Figure 3.**
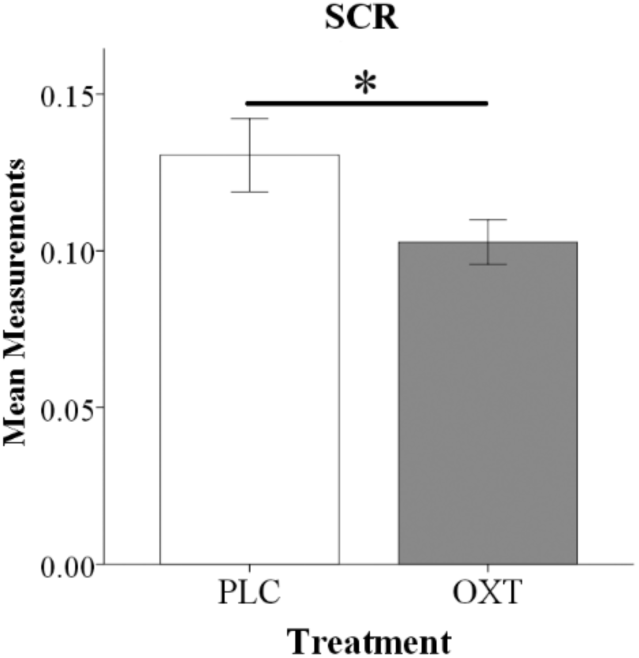
Oxytocin reduced SCR magnitude during the embarrassment task. Histograms show mean SCR magnitude for EE and SE trials combined (* P <0.05)

### fMRI results

For the ROI-based analysis we initially explored whether there were either effects of embarrassment type or gender in the PLC group. This revealed a main effect of embarrassment type in the mPFC (F (1,33) = 5.54, P_FDR_ = 0. 04, η^2^_p_ = 0.14) and in the amygdala (F (1,33) = 7.21, P_FDR_ = 0.01, η^2^_p_ = 0.18) but not in the dAI. This confirmed our expectation that there would be a difference in activation in mentalizing in the SE compared to the EE condition and additionally that there was greater activation in the amygdala during EE compared to SE trials. Examination of OXT by comparing the OXT and the PLC treated groups revealed significant main effect of treatment on right amygdala (F (1,66) = 9.97, P_FDR_ = 0.01, η^2^_p_ = 0.10) and right dAI (F (1,66) = 5.82, P_FDR_ = 0.03, η^2^_p_ = 0.08) responses, with OXT reducing activity in both EE and SE contexts (see Figure 4). No interactions between treatment and gender and embarrassment type were found in these regions. The exploratory whole-brain analysis did not reveal significant main or interaction effects of treatment on the neural level (P_FEW_ < 0.05).

**Figure 4.**
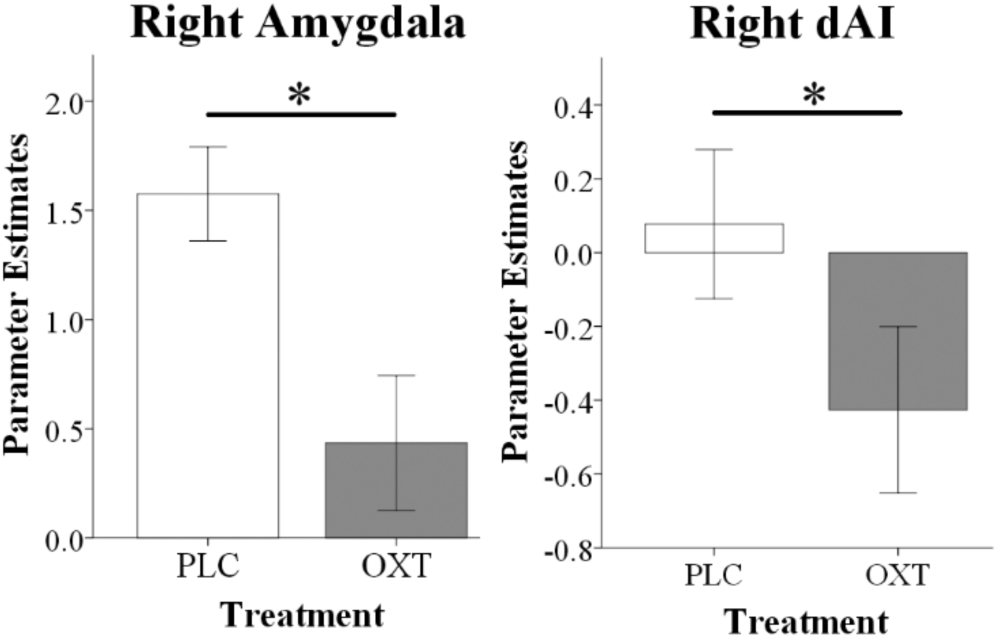
fMRI analysis showing that oxytocin significantly reduced right amygdala and right dAI responses during EE and SE trials. Histograms show parameter estimates for EE and SE trials combined (* P < 0.05)

### Correlation between behavioral, physiological and neural data and trait questionnaires

There was a negative association between overall embarrassment ratings (average of EE and SE) and STAI Trait scores in the OXT but not the PLC group (STAI Trait: PLC, r = 0.06, p = 0.75, OXT, r = -0.34, p = 0.05), suggesting that ratings were particularly increased under OXT in more anxious individuals. However, the correlation difference did not achieve significance (Fisher’s Z Test, z = 1.66, p =0.098). For the neural data there was a similar negative association between right amygdala activation and STAI trait scores in the OXT group but not the PLC group (STAI Trait: PLC, r = 0.13, p = 0.44, OXT, r = -0.35, p = 0.04) and in this case the correlation difference was significant (Fisher’s Z Test, z = 1.99, p = 0.05) (see Figure 5). There was also a negative association between right amygdala activation and the magnitude of the SCR in the OXT group whereas in the PLC group there was a positive association (PLC, r = 0.32, p = 0.12, OXT, r = -0.33, p = 0.06; Fisher’s Z Test, z = 2.40, p = 0.016) (see Figure 5). In contrast, no significant associations between levels of autism (ASQ scores) and behavioral, SCR or neural effects were observed (all ps > 0.15).

**Figure 5.**
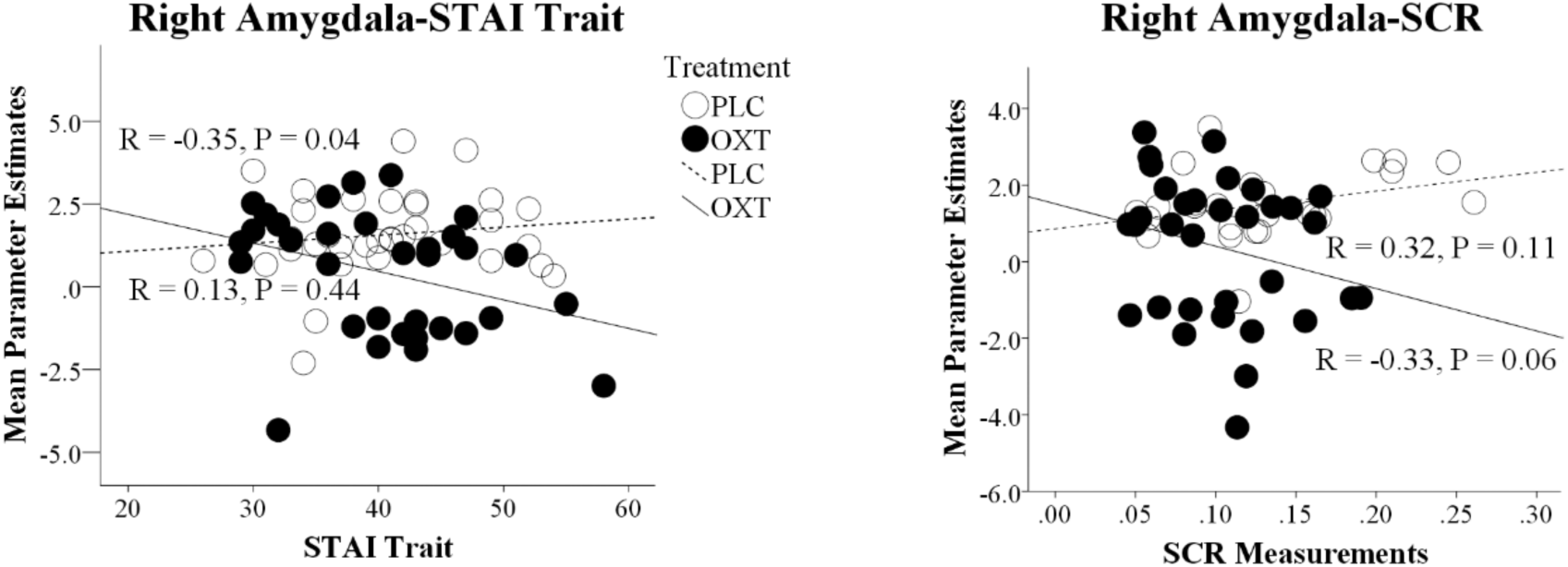
Correlation differences between parameter estimates of right amygdala and trait anxiety scores (STAI Trait) and the magnitude of the skin conductance responses (SCR) in the placebo (PLC) and oxytocin (OXT) groups. Data from EE and SE trials are combined. In both cases the correlation difference between the OXT and PLC groups is significant (Fisher’s Z test p<0.05).

## Discussion

The current experiment demonstrated for the first time that OXT increases both empathic- and self-embarrassment ratings in male and female subjects and that its behavioral effects are associated with decreased responses in the right amygdala and in the right dAI and the magnitude of physiological arousal (SCR). On the other hand, OXT appeared to have no effects on responses in the mentalizing network (mPFC) during either of the embarrassment rating contexts. Furthermore, in contrast with the PLC group the effects of OXT in reducing amygdala responses were negatively correlated with both STAI trait scores and SCR magnitude, implying that it had an anxiolytic effect and particularly in individuals with higher trait anxiety.

Our ROI-based findings in the PLC group that empathic embarrassment primarily increases activation in both mentalizing (mPFC) and emotion processing (amygdala and insula) brain regions is consistent with previous studies (Melchers et al., 2015; Krach et al., 2016; Mueller-Pinzler et al., 2015). Our expectation that the mentalizing component in self-compared to empathic embarrassment would be different was also supported in terms of differential mPFC activation in the PLC group. The effects of OXT in both embarrassment contexts were however restricted to emotion processing regions with reduced amygdala activation, similar to our previous findings for emotional empathy (Geng et al., preprint) and additionally decreased insula activation. Thus, although a number of studies have reported effects of OXT on both dorsal and ventral mPFC activation in the context of self vs other referential and ownership contexts (Zhao et al., 2015; Zhao et al., 2016) it would appear that this does not occur in self- vs other-embarrassment. However, these previous studies demonstrating OXT effects on the mPFC in self- vs. other processing have shown that in both behavioral and neural term’s it reduced the normal self-bias. Thus, it is possible that OXT had no effects on the mPFC since EE and SE ratings were similar and there was therefore no indication of self-bias for embarrassment experience in the present paradigm.

Both EE and SE conditions increased amygdala responses in the PLC group. Although EE trials produced stronger amygdala responses than SE ones following PLC, OXT suppressed them equivalently in both conditions. Several previous studies consistently reported that OXT reduces amygdala reactivity in response to negative emotional stimuli (Kendrick et al., 2017; Kirsch et al., 2005; Striepens et al., 2012). Traditionally this observation has been interpreted as evidence for an anxiolytic effect of OXT, although reduced responses to positive valence stimuli have also been found (Domes et al., 2007). In the current study this anxiolytic effect of OXT is further supported by its reduction of physiological arousal (SCR). There was also a negative association between amygdala activation and trait anxiety scores and the magnitude of the SCR in the OXT group suggesting that it reduced anxiety and particularly in more anxious individuals. This contrasts with our previous findings using the MET where OXT reduced amygdala responses in emotional empathy contexts and this was associated with both increased SCR magnitude and increased empathy ratings (Geng et al., preprint). It seems paradoxical therefore that in the context or empathic and self-embarrassment OXT-reduced amygdala responses were associated with a decreased SCR and yet ratings were still increased. This demonstrates that OXT-induced reduction of amygdala activity per se is not sufficient to infer anxiolytic effects of OXT, and indeed another study has also reported that OXT can promote an anxiogenic response in the context of concomitantly reduced amygdala activation (Riem et al., 2011). In the present study the anxiolytic effects of OXT may have reduced the aversive and threatening affect evoked by the embarrassment stimuli and facilitated a more cognitive appraisal of the social contexts and implications of the situation, thus enhancing embarrassment levels across both, situations experienced either by others or self.

The above anxiolytic interpretation of the effect of OXT is supported by our insula findings where OXT similarly reduced activity in both embarrassment contexts. The anterior insula shows enhanced activation in response to embarrassment (Melchers et al., 2015; Krach et al., 2016; Mueller-Pinzler et al., 2015) and also during perception of others undergoing social or physical pain (Fan et al., 2011; Engen & Singer, 2013). The majority of studies which have found effects of OXT on anterior insula activity have also observed increased responses to negative emotional stimuli consistent with a more anxiogenic effect (Striepens et al., 2012; Yao et al., 2016, Riem et al., 2011; Wigton et al., 2015). Only one other study has reported that OXT reduced anterior insula activity in the context of observing pain in others (Bos et al., 2015). Observing others experiencing embarrassment, which occurred in both EE and SE conditions in our experiment, could be considered as painful emotionally and indeed another study has shown equivalent increased activation in subjects observing physical and social pain (Krach et al., 2016). Thus, OXT may have reduced the feelings of both pain and anxiety experienced during embarrassment, leading to a more cognitive assessment such that subjects rated the level of embarrassment experienced as higher. In this respect it should be noted that subjects in our task were asked to rate the level of embarrassment being experienced in a particular situation and not specifically to rate the intensity of their feelings for others in embarrassing situations, or of their own personal feelings in the self-condition. It is possible therefore that in the context of moral self-conscious emotions such as embarrassment OXT may actually reduce emotional empathy towards others in order to promote a more accurate cognitive assessment of what someone may actually be feeling. This might in turn promote more efficient avoidance of potential embarrassing situations in the future. Further experiments are required to disentangle these cognitive and emotional factors.

In our previous experiment using the MET task we observed that the OXT enhancement of emotional empathy ratings and reduced amygdala activity showed some associations with trait autism, however in the current study we found no such associations. Previous studies have reported decreased responses in the anterior insula to social pain in adults with autism spectrum disorder (Krach et al., 2016) although not for empathic neural responses towards physical pain (Hadjikhani et al., 2014). However, we found no associations between neural responses to embarrassment and autistic traits in healthy subjects. On the other hand, we did find a negative association between trait anxiety and the effect of OXT on the amygdala and SCR, which contrasted with no such association in the PLC group.

This suggested that OXT was particularly attenuating neural and behavioral indices of anxious arousal in subjects with high trait anxiety and resonates with previous findings suggesting that OXT particularly reduced negative appraisal following social stress induction in high trait anxiety subjects (Alvares et al., 2012). These observations conflict with a recent report on OXT-induced increased startle responsivity in high anxious subjects (Schumacher et al., 2018). However, in this previous study OXT specifically increased startle responsivity towards non-social stimuli, further emphasizing how complex interactions between personal and social contextual factors may moderate the specific effects of OXT.

No sex-differential effects were observed in the behavioral effects of OXT on embarrassment ratings or on right amygdala or right insula responses which is consistent with our previous observations that OXT enhanced emotional empathy in the MET (Geng et al., preprint) for both, male and female participants. These findings contrast with other studies reporting sex-dependent responses in amygdala and insula (Gao et al., 2016; Luo et al., 2017) in the context of the impact of positive and negative personal characteristics on face attraction and sub-liminal processing of emotional faces. However, this may reflect the fact that exhibiting empathic-embarrassment and self-embarrassment is of equal adaptive importance in males and females and that OXT only tends to promote or amplify sex-differences in behavioral and neural responses associated with sex-specific priorities in social salience and social preference processing (see Gao et al., 2016).

Findings of the present study need to be interpreted in the context of limitations. Firstly, compared to previous studies that determined sex-differential neural effects of OXT during evaluation social-emotional stimuli the present sample was smaller (n = 70, previous studies enrolled slightly higher samples between arround n = 80-90 subjects, Gao et al., 2016 Luo et al., 2017). The lack of sex-differential effects of OXT on embarrassment therefore needs to be replicated in larger samples. Secondly, our task did not adequately separate cognitive from emotional components of embarrassment in order to provide a better understanding of how OXT attenuation of both amygdala and insula responses resulted in increased ratings of embarrassment.

Overall therefore we have shown for the first time that OXT can enhance ratings of both empathic and self-embarrassment in both males and females, showing that it also influences moral, self-conscious emotional responses. These behavioral effects of OXT are associated with decreased physiological arousal and decreased responses in both the right amygdala and anterior insula but not in mentalizing networks (mPFC). Furthermore, OXT effects on the amygdala are strongest in individuals with high trait anxiety. Thus, OXT in this context may be promoting an anxiolytic effect resulting in a more cognitive rather than emotional appraisal of embarrassment levels.

## Acknowledgements

This study was supported by the National Natural Science Foundation of China (NSFC) grant 563 (grants 31530032; 91632117), the Fundamental Research Funds for the Central Universities of China (ZYGX2015Z002) and the Sichuan Science and Technology Department (2018JY0001).

## Author Contributions Statement

YG, RH and KMK designed this experiment, YG collected the data, YG, WZ, FZ, KMK and BB analyzed the data, YG, WZ, XM, SY, RH, BB and KMK interpreted the results. YG, BB and KK wrote the paper.

## Conflicts of Interest Statement

Authors declare no conflicts of interest.

## Supplementary table 1

**Table S1.** Demographics of ages and questionnaire scores for study subjects in current experiment (mean ± SEM)

| Measurements | PLC | OXT | T-value | P |
| --- | --- | --- | --- | --- |
| Age (yr) | 21.86 ± 0.33 | 22.03 ± 0.36 | -0.35 | 0.73 |
| Education | 15.89 ± 0.28 | 16.09 ± 0.28 | -0.51 | 0.61 |
| Empathy Quotient (EQ) | 38.17 ± 1.72 | 37.83 ± 1.36 | 0.16 | 0.88 |
| Beck Depression Inventory (BDI) | 7.43 ± 1.05 | 4.91 ± 0.89 | 1.83 | 0.07 |
| State and Trait Anxiety Inventory (STAI)_State | 39.74 ± 1.54 | 37.94 ± 1.14 | 0.94 | 0.35 |
| State and Trait Anxiety Inventory (STAI)_Trait | 40.63 ± 1.19 | 40.37 ± 1.25 | 0.15 | 0.88 |
| Positive and Negative Affect Scale (PANAS)_Positive | 29.97 ± 0.99 | 29.17 ± 0.88 | 0.61 | 0.55 |
| Positive and Negative Affect Scale (PANAS)_Negative | 19.91 ± 1.16 | 18.37 ± 0.90 | 1.05 | 0.30 |
| The Adult Autism Spectrum Quotient (ASQ) | 20.14 ± 0.98 | 19.06 ± 1.13 | 0.72 | 0.47 |
| Wong Law Emotional Intelligence Scale-Chinese(WLEIS-C) | 83.71 ± 1.78 | 83.00 ± 2.33 | 0.24 | 0.81 |
| Liebowitz Social Anxiety Scale (LSAS)_Fear | 43.66 ± 1.82 | 42.74 ± 1.77 | 0.36 | 0.72 |
| Liebowitz Social Anxiety Scale (LSAS)_Avoidance | 42.09 ± 1.87 | 40.83 ± 1.86 | 0.48 | 0.64 |
| The Self-Esteem Scale (SES) | 31.71 ± 0.63 | 32.60 ± 0.67 | -0.96 | 0.34 |

## References

Ali F, Amorim IS, Chamorro-Premuzic T. Empathy deficits and trait emotional intelligence in psychopathy and Machiavellianism. Pers Individ Dif (2009) 47:758–762. doi:10.1016/j.paid.2009.06.016

Alvares GA, Chen NT, Balleine BW, Hickie IB, Guastella AJ. Oxytocin selectively moderates negative appraisals in high trait anxious males. Psychoneuroendocrinology (2012) 37: doi:10.1016/j.psyneuen.2012.04.018

Baron-Cohen S, Wheelwright S, Skinner R, Martin J, Clubley E. The autism-spectrum quotient (AQ): evidence from Asperger syndrome/high-functioning autism, males and females, scientists and mathematicians. J Autism Dev Disord (2001) 31:5–17.

Baron-Cohen S, Wheelwright S. The Empathy Quotient (EQ): An investigation of adults with Asperger Syndrome or High Functioning Autism and normal sex differences. J Autism Dev Disord (2004) 34:163–175. doi:10.1023/B:JADD.0000022607.19833.00

Bartz JA, Zaki J, Bolger N, Hollander E, Lduwing NN, Kolevzon A, Ochsner KN. Oxytocin selectively improves empathic accuracy. Psychol Sci (2010) 21: doi:10.1177/0956797610383439

Beck AT. Beck Depression Inventory. Depression (1961) 2006:2–4. doi:10.1093/ndt/gfr086

Bos PA, Montoya ER, Hermans EJ, Keysers C, van Honk J. Oxytocin reduces neural activity in the pain circuitry when seeing pain in others. Neuroimage (2015) 113:217–224.

Brett M, Anton J-L, Valabregue R, Poline J-B. Region of interest analysis using the MarsBar toolbox for SPM 99. Neuroimage (2002) 16: S497.

Domes G, Heinrichs M, Glascher J, Buchel C, Braus DF, Herpertz SC. Oxytocin attenuates amygdala responses to emotional faces regardless of valence. Biol Psychiatry (2007) 62:1187–1190. doi:10.1016/j.biopsych.2007.03.025

Domes G, Heinrichs M, Michel A, Berger C, Herpertz SC. Oxytocin improves “mind-reading” in humans. Biol Psychiatry (2007) 61:731–733.

Engen HG, Singer T. Empathy circuits. Curr Opin Neurobiol (2013) 23: doi:10.1016/j.conb.2012.11.003

Fan Y, Duncan NW, de Greck M, Northoff G. Is there a core neural network in empathy? An fMRI based quantitative meta-analysis. Neurosci Biobehav Rev (2011) 35:903–911. doi:10.1016/j.neubiorev.2010.10.009

Feeser M, Fan Y, Weigand A, Hahn A, Gärtner M, Böker H, Grimm S, Bajbouj M. Oxytocin improves mentalizing–pronounced effects for individuals with attenuated ability to empathize. Psychoneuroendocrinology (2015) 53:223–232.

Gamer M, Zurowski B, Buchel C. Different amygdala subregions mediate valence-related and attentional effects of oxytocin in humans. Proc Natl Acad Sci U S A (2010) 107:9400–9405. doi:10.1073/pnas.1000985107

Gao S, Becker B, Luo L, Geng Y, Zhao W, Yin Y, Hu J, Gao Z, Gong Q, Hurlemann R, et al. Oxytocin, the peptide that bonds the sexes also divides them. Proc Natl Acad Sci U S A (2016) 113:7650–7654. doi:10.1073/pnas.1602620113

Geng Y, Zhao W, Zhou F, Ma X, Yao S, Hurlemann R, Becker B, Kendrick K. Oxytocin enhancement of emotional empathy: generalization across cultures and effects on amygdala activity. bioRxiv (2018) Preprint available at: http://biorxiv.org/content/early/2018/04/26/307256.abstract

Guastella AJ, Hickie IB, McGuinness MM, Otis M, Woods EA, Disinger HM, Chan HK, Chen TF, Banati RB. Recommendations for the standardisation of oxytocin nasal administration and guidelines for its reporting in human research. Psychoneuroendocrinology (2013) 38:612–625. doi:10.1016/j.psyneuen.2012.11.019

Hadjikhani N, Zurcher NR, Rogier O, Hippolyte L, Lemonnier E, Ruest T, Ward N, Lasalle A, Gillberg N, Nillstedt E, Helles A, Gilberg C, Solomon P, Prkachin KM, Gillberg C. Emotional contagion for pain is intact in autism spectrum disorders. Transl Psychiatry (2014) 4: doi:10.1038/tp.2013.113.

Hubble K, Daughters K, Manstead A, Rees A, Thapar A, Van SG. Oxytocin Increases Attention to the Eyes and Selectively Enhances Self-Reported Affective Empathy for Fear. Neuropsychologia (2017) 106:350.

Hurlemann R, Patin A, Onur OA, Cohen MX, Baumgartner T, Metzler S, Dziobek I, Gallinat J, Wagner M, Maier W, et al. Oxytocin Enhances Amygdala-Dependent, Socially Reinforced Learning and Emotional Empathy in Humans. J Neurosci (2010) 30:4999–5007.

Kendrick KM, Guastella AJ, Becker B. (2017) Overview of Human Oxytocin Research, in Curr Top Behav Neurosci, doi:10.1007/7854_2017_19

Kirsch P. Oxytocin Modulates Neural Circuitry for Social Cognition and Fear in Humans. J Neurosci (2005) 25:11489–11493. doi:10.1523/JNEUROSCI.3984-05.2005

Krach S, Müller-Pinzler L, Rademacher L, Sören D, Frieder S ·, Paulus M. Neuronal pathways of embarrassment. (2016) 7:37–42. doi:10.1007/s13295-016-0024-4

Liebowitz MR. Liebowitz Social Anxiety Scale. Mod Probl Pharmapsychiatry (1987) 22:141–73.

Lombardo M V., Barnes JL, Wheelwright SJ, Baron-Cohen S. Self-referential cognition and empathy in autism. PLoS One (2007) 2: doi:10.1371/journal.pone.0000883

Luo L, Becker B, Geng Y, Gao S, Zhao W, Yao S, Zheng X, Ma X, Gao Z, Hu J, Kendrick KM. Sex-dependent neural effect of oxytocin during subliminal processing og negative emotional faces. Neuroimage (2017) 162: doi:10.1016/j.neuroimage.2017.08.079

Melchers M, Markett S, Montag C, Trautner P, Weber B, Lachmann B, Buss P, Heinen R, Reuter M. Reality TV and vicarious embarrassment: An fMRI study. Neuroimage (2015) 109:109–117. doi:10.1016/j.neuroimage.2015.01.022

Miller RS. Empathic embarrassment: Situational and personal determinants of reactions to the embarrassment of another. J Pers Soc Psychol (1987) 53:1061–1069. doi:10.1037/0022-3 514.53.6.1061

Moya-Albiol L, Herrero N, Bernal MC. The neural bases of empathy. Rev Neurol (2010) 50:89–100. doi:rn2009111[pii]

Müller-Pinzler L, Gazzola V, Keysers C, Sommer J, Jansen A, Frässle S, Einhäuser W, Paulus FM, Krach S. Neural pathways of embarrassment and their modulation by social anxiety. Neuroimage (2015) doi:10.1016/j.neuroimage.2015.06.036

Olff M, Frijling JL, Kubzansky LD, Bradley B, Ellenbogen MA, Cardoso C, Bartz JA, Yee JR, van Zuiden. The role of oxytocin in social bonding, stress regulation and mental health: an update on moderating effects of context and interindividual differences. Psychoneuroendocrinology (2013) 38: doi:10.1016/j.psyneuen.2013.06.019

Preston SD, Bechara A, Damasio H, Grabowski TJ, Stansfield RB, Mehta S, Damasio AR. The neural substrates of cognitive empathy. Soc Neurosci (2007) 2:254–275. doi:10.1080/17470910701376902

Radke S, de Bruijn ERA. Does oxytocin affect mind-reading? A replication study. Psychoneuroendocrinology (2015) 60:75–81. doi:10.1016/j.psyneuen.2015.06.006

Riem MME, Bakermans-Kranenburg MJ, Pieper S, Tops M, Boksem MAS, Vermeiren RRJM, van IJzendoorn MH, Rombouts SARB. Oxytocin Modulates Amygdala, Insula, and Inferior Frontal Gyrus Responses to Infant Crying: A Randomized Controlled Trial. Biol Psychiatry (2011) 70:291–297. doi:10.1016/j.biopsych.2011.02.006

Riem MME, Bakermans-Kranenburg MJ, Voorthuis A, van IJzendoorn MH. Oxytocin effects on mind-reading are moderated by experiences of maternal love withdrawal: An fMRI study. Prog Neuropsychopharmacol Biol Psychiatry (2014) 51:105–112.

Scheele D, Kendrick KM, Khouri C, Kretzer E, Schlapfer TE, Stoffel-Wagner B, Gunturkun O, Maier W, Hurlemann R. An oxytocin-induced facilitation of neural and emotional responses to social touch correlates inversely with autism traits. Neuropsychopharmacology (2014) 39:2078–2085. doi:10.1038/npp.2014.78

Schumacher S, Oe M, Wilhelm FH, Rufer M, Heinrichs M, Weidt S, Morgeli H, Martin-Soelch. Does trait anxiety influence effects of oxytocin on eye-blink startle reactivity? A randomized, double-blind, placebo-controlled crossover study. PLoS One (2018) 13: doi:10.1371/journal.pone.0190809.

Shamay-Tsoory SG. The neural bases for empathy. Neuroscientist (2011) 17:18–24. doi:10.1177/1073858410379268

Spielberger CD, Gorsuch RL, Lushene RE. Manual for the state-trait anxiety inventory. (1970)

Stern RM, Ray WJ, Quigley KS. Psychophysiological recording. Oxford University Press, USA (2001).

Striepens N, Kendrick KM, Maier W, Hurlemann R. Prosocial effects of oxytocin and clinical evidence for its therapeutic potential. Front Neuroendocrinol (2011) 32:426–450. doi:10.1016/j.yfrne.2011.07.001

Striepens N, Scheele D, Kendrick KM, Becker B, Schafer L, Schwalba K, Reul J, Maier W, Hurlemann R. Oxytocin facilitates protective responses to aversive social stimuli in males. Proc Natl Acad Sci U S A (2012) 109:18144–18149. doi:10.1073/pnas.1208852109

Tully EC, Ames AM, Garcia SE, Donohue MR ose. Quadratic Associations Between Empathy and Depression as Moderated by Emotion Dysregulation. J Psychol (2016) 150:15–35. doi:10.1080/00223980.2014.992382

Tzourio-Mazoyer N, Landeau B, Papathanassiou D, Crivello F, Etard O, Delcroix N, Mazoyer B, Joliot M. Automated anatomical labeling of activations in SPM using a macroscopic anatomical parcellation of the MNI MRI single-subject brain. Neuroimage (2002) 15:273–289. doi:10.1006/nimg.2001.0978

Watson D, Clark LA, Tellegen A. Development and validation of brief measures of positive and negative affect: the PANAS scales. J Pers Soc Psychol (1988) 54:1063–1070. Available at: https://www.ncbi.nlm.nih.gov/pubmed/3397865

Wigton R, Radua J, Allen P, Averbeck B, Meyer-Lindenberg A, McGuire P, Shergill SS, Fusar-Poli P. Neurophysiological effects of acute oxytocin administration: systematic review and meta-analysis of placebo-controlled imaging studies. J Psychiatry Neurosci (2015) 40: doi:10.1503/jpn.130289

Wong C-S, Law KS. The effects of leader and follower emotional intelligence on performance and attitude: An exploratory study. Leadersh Q (2002) 13:243–274.

Yao S, Becker B, Zhao W, Zhao Z, Kou J, Ma X, Geng Y, Ren P, Kendrick KM. Oxytocin Modulates Attention Switching Between Interoceptive Signals and External Social Cues. Neuropsychopharmacology (2017) doi:10.1038/npp.2017.189

Zhao W, Geng Y, Luo L, Zhao Z, Ma X, Xu L, Yao S, Kendrick KM. Oxytocin Increases the Perceived Value of Both Self- and Other-Owned Items and Alters

Medial Prefrontal Cortex Activity in an Endowment Task. Front Hum Neurosci (2017) 11: doi:10.3389/fnhum.2017.00272

Zhao W, Yao S, Li Q, Geng Y, Ma X, Luo L, Xu L, Kendrick KM. Oxytocin blurs the self-other distinction during trait judgments and reduces medial prefrontal cortex responses. Hum Brain Mapp (2016) 37:2512–2527. doi:10.1002/hbm.23190

